# Endophytic microbiome variation at the level of a single plant seed

**DOI:** 10.1101/2021.04.27.441449

**Authors:** AF Bintarti, A Sulesky-Grieb, N Stopnisek, A Shade

## Abstract

Like other plant compartments, the seed harbors a microbiome. The members of the seed microbiome are the first to colonize a germinating seedling, and they initiate the trajectory of microbiome assembly for the next plant generation. Therefore, the members of the seed microbiome are important for the dynamics of plant microbiome assembly and the vertical transmission of potentially beneficial symbionts. However, it remains challenging to assess the microbiome at the individual seed level (and, therefore, for the future individual plant) due to low endophytic microbial biomass, seed exudates that can select for particular members, and high plant and plastid contamination of resulting reads. Here, we report a protocol for extracting metagenomic DNA from an individual seed (common bean, *Phaseolus vulgaris* L.) with minimal disruption of host tissue, which we expect to be generalizable to other medium-and large-seed plant species. We applied this protocol to quantify the 16S rRNA V4 and ITS2 amplicon composition and variability for individual seeds harvested from replicate common bean plants grown under standard, controlled conditions to maintain health. Using metagenomic DNA extractions from individual seeds, we compared seed-to-seed, pod-to-pod, and plant-to-plant microbiomes, and found highest microbiome variability at the plant level. This suggests that several seeds from the same plant could be pooled for microbiome assessment, given experimental designs that apply treatments at the maternal plant level. This study adds protocols and insights to the growing toolkit of approaches to understand the plant-microbiome engagements that support the health of agricultural and environmental ecosystems.

## Introduction

Seed microbiomes offer a reservoir of microbiota that can be vertically passed from maternal plants to offspring (Mitter et al. 2017; Shade et al. 2017; Truyens et al. 2015) and some of these members have plant-beneficial phenotypes (Adam et al. 2018; Berg and Raaijmakers 2018; Bergna et al. 2018; López-López et al. 2010). Therefore, the seed microbiome is expected to a play a key role in plant health and fitness (Barret et al. 2015), and especially in the assembly and establishment of the developing plant’s microbiome (Chesneau et al. 2020). This expected importance of the seed microbiome has fueled recent research activity to use high-throughput sequencing to characterize the seed microbiomes of various plants (e.g., Chartrel et al. 2021; Dai et al. 2020; Eyre et al. 2019; Raj et al. 2019; Rodríguez et al. 2020; Xing et al. 2018).

Seed microbiomes include microbial members that live on the seed surface as epiphytes and members that colonize inside the internal tissue of the seed as endophytes (Nelson 2018). Among these microbiome members, endophytes that closely associate with endosperm and embryo are more likely to be transmitted to the next plant generations than are seed-associated epiphytes (Barret et al. 2016; Nelson 2018). By itself, an endophytic association does not confirm that there is a functional benefit or co-evolutionary relationship between the plant and the microbiome member (Nelson 2018). However, endophytic microbes offer the first source of inoculum for the germinating seedling (as reviewed in Nelson 2018; Vujanovic and Germida 2017), and, given the potential for priority effects or pathogen exclusion, these members can have implications for the mature plant’s microbial community structure. Therefore, understanding the endophytic seed microbiome is expected to provide insights into mechanisms of seed facilitation of microbiome assembly and the vertical transmission of microbiome members over plant generations.

As is true for other plant compartments, different plant species or divergent crop lines/varieties/cultivars often have different seed microbiome composition or structure (Wassermann et al. 2019; Klaedtke et al. 2016; Johnston-Monje and Raizada 2011; López-López et al. 2010). However, many seed microbiome studies have reported generally high variability across seed samples from the same plant type and treatment (Bergna et al. 2018; López-López et al. 2010; Bintarti et al. 2020), with strong explanatory value of either seed origin/seed lot, geographic region or soil edaphic conditions (Chartrel et al. 2021; Klaedtke et al. 2016; Johnston-Monje and Raizada 2011; but see also Adam et al. 2018 for an exception). While these insights may call into question the proportion of “inherited” versus acquired seed microbiome members, the high microbiome variability may be in part due to methods applied to extract the microbial DNA from the seed compartment, and different methods applied across studies. For instance, some studies surface sterilize the seeds while others do not; some germinate the seed prior to microbiome analysis while others do not, etc. One source of microbiome variability could be the common practice of the pooling of many seeds from the same or different plants to produce a composite seed microbiome sample for DNA extraction. Because multiple seeds are investigated at once, it is unclear at what level the most microbiome variability is highest– the seed, the pod or fruit, the plant, or the field or treatment. This information is required to determine the necessary sample size in well-powered experimental designs. More importantly, the question of vertical transmission cannot directly be addressed without seed microbiome assessment of an individual.

Our study objectives were: 1) to determine the appropriate observational unit of endophytic seed microbiome assessment for common bean (*Phaseolus vulgaris* L) by quantifying seed-to-seed, pod-to-pod, and plant-to-plant variability in 16S rRNA V4 and ITS2 amplicon analyses; and 2) to develop a robust protocol for individual seed microbiome extraction that could be generally applied to other plants that have similarly medium-to large-sized seeds. We found that that plant-to-plant variability under controlled growth conditions exceeded within-plant variability and conclude that seeds can be pooled by maternal plant (but, not across different plants) in study designs that aim to compare seed microbiomes resulting from treatments applied at the plant level.

## Materials and Methods

### Plant growth conditions

Because we targeted the endophytic seed microbiome, surface sterilization of the bean seeds was conducted before germination and planting. To sterilize, seeds were soaked in a solution of 10% bleach with 0.1% Tween20 for 15 minutes, then rinsed four times with sterile water. The final rinse water was plated on tryptic soy agar (TSA) and potato dextrose agar (PDA) plates to test for sterilization efficacy. Sterilized seeds were placed in Petri dishes on sterile tissue paper moistened with sterile water, and allowed to germinate in in the dark for four days. After four days, the radicle had emerged and the germinated seeds were ready to be transferred to the growth chamber. The germinated seeds were planted in three 4.54 L (1-gallon) pots filled with a 50:50 v/v mixture of agricultural bean field soil and vermiculite. The pots were placed in a BioChambers model SPC-37 growth chamber with a 14-h day/10-h night cycle at 26°C and 22°C, respectively, 260 mE light intensity, and 50% relative humidity. All plants received 300 mL of water every other day and 200 mL of half-strength Hoagland solution (Hoagland and Arnon 1950) once a week.

### Study design

We planted three germinated seeds per pot and culled to one seedling per pot at the early vegetative growth stage. There were three plant replicates designated as A, B, and C, grown under the above-described conditions for normal, healthy growth. The three plants produced different numbers of pods and total seeds (plant A = 5 pods, 22 seeds; plant B = 6 pods, 29 seeds; and plant C = 7 pods, 26 seeds) with the number of seeds varying across pods (2 to 6 seeds per pod). We aimed to balance and maximize number of seeds across plants. Therefore, we extracted metagenomic DNA from 3 pods from plants A and C, and 6 pods from plant B, with 3 to 4 seeds in each pod. For the 16S V4 analysis we had 3 pods from plant A (A1, A2 and A3= 4 seeds), 6 pods from plant B (B1 through B6 = 4 seeds), and 3 pods from plant C (C5= 3 seeds, C6 and C7= 4 seeds) for a total of 47 individual seed samples analyzed. For the ITS2 analysis, we were unable to amplify fungal target DNA from pod A1 or pod B1, for a total of 45 individual seed samples analyzed.

### Seed harvest and endophyte metagenomic DNA extraction

Once the plants reached maturity at the R9 growth stage (yellowing leaves and dry pods), the seeds were harvested for endophytic microbiome analysis. Seeds were distinguished by plant and pod. The endophytic microbiome of each seed was extracted and sequenced individually. To extract the endophytic metagenomic DNA (mgDNA), a protocol was adapted from Barret et al. 2015 and Rezki et al. 2018. First, the seeds were surface-sterilized as above and the seed coat was carefully removed using sterilized forceps. Each seed was then soaked in 3 mL of PBS solution with 0.05% Tween20 (hereafter, “soaking solution”) overnight at 4°C with constant agitation of 170 rpm. Since low levels of microbial biomass are expected in single seed extractions, positive and negative controls were included in the extraction protocol. This ensures that if no extractable microbial DNA is present in a sample that it is representative of the sample, rather than the extraction methods. A mock community was used as a DNA extraction positive control by adding one, 75 µL aliquot of the ZymoBIOMICS™ Microbial Community Standard (Zymo Research, Irvine, CA, United States) to 3 mL of the soaking solution immediately prior to conducting the extraction protocol. Sterile soaking solution (3 mL) was used as a negative DNA extraction control.

After soaking overnight, the samples were centrifuged at 4500xg for 60 minutes at 4°C to pellet any material that had been released from the seed tissues. After centrifugation the seed was removed, and the pelleted material was resuspended in 1-2 mL of supernatant and transferred to a microcentrifuge tube for DNA extraction using the E.Z.N.A^®^ Bacterial DNA Kit (Omega Bio-tek, Inc. Norcross, GA, United States). The manufacturer’s Centrifugation Protocol was used with minor modifications. Specifically, the pelleted seed material was suspended in TE buffer (step 4), the incubation for the lysozyme step was extended to 20 minutes, 30 µL of elution buffer was used, and the elution step was extended to a 15 minute incubation. These modifications were performed to maximally recover the limited amount of mgDNA expected from a single seed. We detail the standard operating protocol, and provide notes on the alternatives that we tested in optimizing this protocol in the Supplementary Material.

### PCR amplification and amplicon sequencing

To confirm successful DNA extraction from the seed pellet, DNA quantification and target gene polymerase chain reaction (PCR) assays were performed. First, the DNA extracted from the seed samples and the positive and negative controls were quantified using the Qubit™dsDNA BR Assay Kit (ThermoFisher Scientific, Waltham, MA, United States). Then, PCR amplification and sequencing of the V4 region of 16S rRNA bacterial/archaeal gene and the ITS2 region of the ITS fungal gene were performed. The V4 region of 16S rRNA gene amplification was conducted using 515f (5’-GTGCCAGCMGCCGCGGTAA-3’) and 806r (5’-GGACTACHVGGGTWTCTAAT-3’) universal primers (Caporaso et al. 2011) under the following conditions: 94°C for 3 min, followed by 35 cycles of 94°C (45 s), 50°C (60 s), and 72°C (90 s), with a final extension at 72°C (10 min). The amplification was performed in 25 µl mixtures containing 12.5 µl GoTaq® Green Master Mix (Promega, Madison, WI, United States), 0.625 µl of each primer (20 µM), 2 µl of DNA template (∼1 ng per µl), and 9.25 µl nuclease free water. The mgDNA (concentration of ∼ 1 ng per µl) was sequenced at the Research Technology Support Facility (RTSF) Genomics Core, Michigan State sequencing facility using the Illumina MiSeq platform v2 Standard flow cell. The sequencing was performed in a 2×250bp paired end format.

The PCR amplification of the ITS2 region of the ITS gene was performed using ITS86f (5’-GTGAATCATCGAATCTTTGAA-3′) and ITS4 (5’-TCCTCCGCTTATTGATATGC-3’) primers (Op De Beeck et al. 2014) with addition of index adapters by the RTSF Genomics Core. The PCR amplification of the ITS2 was conducted under the following conditions: 95°C for 2 min, followed by 40 cycles of 95°C (30 s), 55°C for (30 s), and 72°C for (1 min), with a final extension at 72°C for 10 min. The amplification was performed in 50 µl mixture containing 20 µl GoTaq®Green Master Mix (Promega, Madison, WI, United States), 1 µl of each primer (10 µM), 1 µl of DNA template (∼ 1 ng per µl), and 27 µl nuclease free water. The PCR products were purified using QIAquick® PCR Purification Kit (QIAGEN, Hilden, Germany). Purified PCR products with a concentration range 6-10 ng per µl were sequenced at the RTSF Genomics Core using Illumina MiSeq platform v2 Standard flow cell and 2×250bp paired end format.

### Sequence analysis

The USEARCH pipeline (v.10.0.240) was used to merge paired-end bacterial/archaeal raw reads, filter for low-quality sequences, dereplicate, remove singletons, denoise, and check for chimeras (Edgar and Flyvbjerg 2015). An in-house open reference strategy was performed for OTU clustering (Rideout et al. 2014). First, closed-reference OTU picking was performed by clustering the quality filtered reads against the SILVA database (v.132) (Quast et al. 2013) at 97% identity using USEARCH algorithm (usearch_global command) (Edgar 2010). Then, de novo OTU picking process was performed on the reads that failed to match the reference using UPARSE-OTU algorithm (cluster_otus command) (Edgar 2013) at 97% identity. Finally, closed-reference and de novo OTUs were combined into a full set of representative sequences. The merged sequences were then mapped back to the representative sequences using the usearch_global command.

Sequence alignment, taxonomy assignment, non-bacteria/archaea filtering, and phylogenetic diversity calculation were performed using QIIME 1.9.1. The representative sequences were aligned against the SILVA database (v.132) (Caporaso, Kuczynski, et al. 2010) using PyNAST (Caporaso, Bittinger, et al. 2010). The unaligned OTUs and sequences were excluded from the OTU table and the representative sequences file, respectively. Taxonomy assignment was performed using the default classifier method (UCLUST algorithm) at a minimum confidence of 0.9 (Edgar 2010) using SILVA database (v.132) as the reference. Plant contaminants (chloroplast and mitochondria) and unassigned taxa were removed from the OTU table and the representative sequences using filter_taxa_from_otu_table.py and filter_fasta.py command. Filtering the microbial contaminants from the OTU table was conducted in R (v.3.4.2) (R Core Development Team) using the microDecon package (McKnight et al. 2019). Reads were normalized using Cumulative Sum Scaling (CSS) method in metagenomeSeq Bioconductor package on R (Paulson et al. 2013).

The fungal ITS raw reads were processed using the USEARCH (v.10.0.240) pipeline. Read processing included merging paired-end reads, removing primers using cutadapt (v.2.1) (Martin 2011), dereplication, and singleton removal. OTUs were picked and chimeras removed using de novo clustering at 97% identity threshold with the UPARSE-OTU algorithm (cluster_otus command, Edgar 2013). Then, all merged sequences were mapped to the clustered reads using usearch_global command to generate an OTU table. Fungal taxonomic classification was performed in CONSTAX (Gdanetz et al. 2017) using RDP Classifier (v.11.5) (Cole et al. 2014; Wang et al. 2007) at a minimum confidence of 0.8 and with the UNITE reference database (release 01-12-2017). Plant and microbial contaminants removal and read normalization were performed in R (v.3.4.2). Plant contaminants were removed from the OTU table by filtering out OTUs that were assigned into Kingdom Plantae. Microbial contaminants were removed using the microDecon package (McKnight et al. 2019). The CSS method from the metagenomeSeq Bioconductor package was performed to normalize the fungal reads (Paulson et al. 2013).

### Microbial community analysis

Microbiome statistical analyses were conducted in R (v.3.4.2) (R Core Development Team). Microbial alpha and beta diversity were calculated on the CSS-normalized OTU table using the vegan package (v.2.5-7) (Oksanen et al. 2019). Richness and Faith’s phylogenetic diversity were used to analyze the bacterial/archaeal alpha diversity. For fungal alpha diversity, we used richness. The evenness of the seed microbiomes was visualized using rank-abundance curves (Phyloseq package (v.1.28.0) in R (McMurdie and Holmes 2013)). Differences in alpha diversity among plants and pods were determined by fitting the Linear Mixed-Effects Model (LMM) using the lme() function of the nlme package (version 3.1-152) (Pinheiro et al. 2021). We performed LMM because the study has an unbalanced nested design with pod as the random factor, nested within plant as the fixed factor. Microbial composition and relative abundance were analyzed using the Phyloseq package (v.1.28.0) in R (McMurdie and Holmes 2013).

Beta diversity was calculated using Jaccard distances and visualized using principal coordinate analysis (PCoA) plot. We used the Jaccard index, which is based on presence-absence, rather than a metric based on relativized abundance because we reasoned that the seed microbiome members are likely to be dormant inside the seed prior to germination (Cope-Selby et al. 2017), and that any differences in relative abundances are not directly attributablem to competitive fitness outcomes inside the seed. Furthermore, exponential growth would allow that any viable cell successfully packaged and passaged via the seed could, in theory, successfully colonize the new plant. Nested permutational multivariate analysis of variance (PERMANOVA) using the function nested.npmanova() from the BiodiversityR package (Kindt 2020) was performed to assess the microbial community structure among plants and pods. We performed multivariate analysis to check the homogeneity of dispersion (variance) among groups using the function betadisper() (Oksanen et al. 2019). We performed PERMADISP to test the significant differences in dispersions between groups and Tukey’s HSD test to determine which groups differ in relation to the dispersions (variances).

Power analysis and sample size were calculated using the pwr.t.test() function from the pwr package (v.1.3-0). We performed power analysis of two-category t-test. Because the most microbiome variability was observed across plants, we pooled individual seed sequence profiles in silico at the plant level for this analysis. We calculated Cohen’s d effect size given the information of mean and standard deviation of bacterial/archaeal alpha diversity (richness and phylogenetic diversity) from three plant samples from this study: Plant A (n = 12; richness: M = 30.58, SD = 6.42, phylogenetic diversity: M = 4.17, SD = 0.89), Plant B (n = 24; richness: M = 18.21, SD = 7.35, phylogenetic diversity: M = 2.92, SD = 0.82) and Plant C (n= 11; richness: M = 19.09, SD = 10.95, phylogenetic diversity: M = 3.09, SD = 1.39). We calculated the common standard deviation (σpool of all groups) using the above information, then we calculated Cohen’s d effect size for both richness and phylogenetic diversity. Cohen’s d effect size was defined by calculating the difference between the largest and smallest means divided by the square root of the mean square error (or the common standard deviation). Power analysis was run with Hedges’s g effect size (corrected with Cohen’s d effect size) and significant level of 0.05.

### Data and code availability

The computational workflows for sequence processing and ecological statistics are available on GitHub (https://github.com/ShadeLab/Bean_seed_variability_Bintarti_2021). Raw sequence data of bacteria/archaea and fungi have been deposited in the Sequence Read Archive (SRA) NCBI database under Bioproject accession number PRJNA714251.

## Results

### Sequencing summary and microbiome coverage

A total of 5,056,769 16S rRNA V4 and 8,756,009 ITS2 quality reads were generated from 47 mgDNA samples purified from individual seeds for bacteria/archaea, and from 45 samples for fungi. We removed more than 90 % of reads that were plant contaminants (**Fig. S1**), resulting in 17,128 and 67,878 16S rRNA bacterial/archaeal and ITS fungal reads, respectively. After removing plant and microbial contaminants, we determined 211 bacterial/archaeal and 57 fungal operational taxonomic units (OTUs) defined at 97% sequence identity. While the majority of individual seeds from plants A and B had exhaustive to sufficient sequencing effort, some seeds from plant C did not (**Fig. 1a**). However, the fungal rarefaction curves reached asymptote and had sufficient effort (**Fig. 1b**). Both bacterial/archaeal and fungal seed microbiomes were highly uneven with few dominant and many rare taxa, as typical for microbiomes (**Fig 1c,d**).

**Fig 1.**
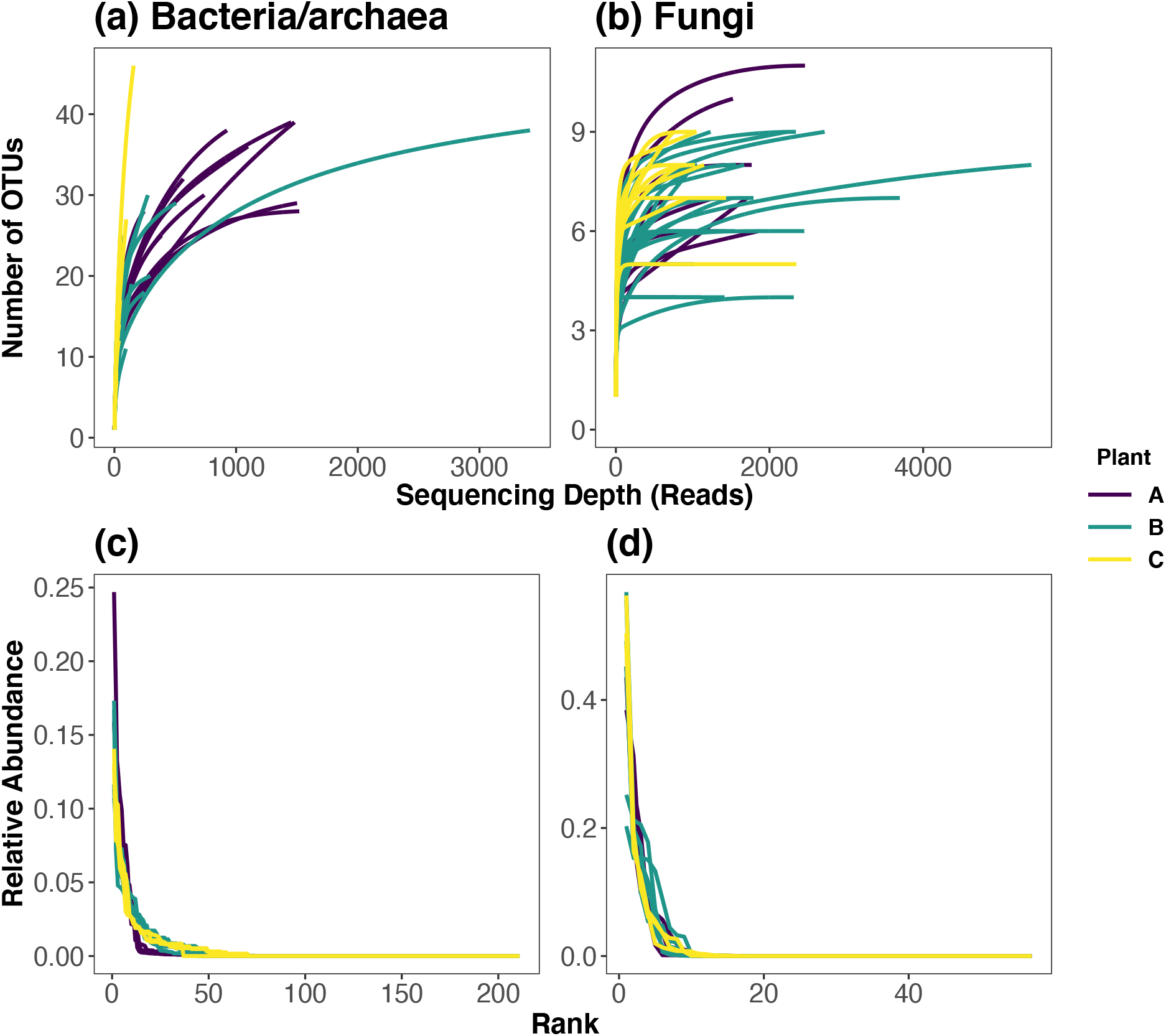
Rarefaction curves of bacteria/archaea (a) and fungi (b) from seed samples (marked) at 97 % of clustering threshold were constructed by plotting the OTU number after decontamination (microbial contaminants removal) to the sequence (read) number. The rarefaction curves were constructed using vegan package (v2.5-4). Rank abundance curve of decontaminated and normalized bacterial/archaeal (c) and fungal (d) OTU tables. Samples (n=47 and n=45 for bacteria/archaea and fungi, respectively) were grouped by plant.

### Microbiome Diversity

There were differences in bacterial/archaeal richness among seeds from different plants (LMM; df = 2, F-value = 6.91, p-value = 0.015) (**Fig. 2a**), where plant B and C had lower seed richness than plant A (Tukey’s HSD post hoc test; p-value = 0.001 and 0.006, respectively). However, bacterial/archaeal richness among seeds from pods collected from the same plant were not different (LMM, p-value > 0.05) (**Fig. 2b**). Similarly, bacterial/archaeal phylogenetic diversity were different among seeds collected from different plants (LMMs; df = 2, F-value = 6.56, p-value = 0.003) (**Fig. 2c**), but not among seeds from pods within the same plant (LMM, p-value > 0.05) (**Fig. 2d**). Plants B and C had lower seed microbiome bacterial/archaeal phylogenetic diversity compared to plant A (Tukey’s HSD post hoc test, p-value = 0.001 and 0.013, respectively). We observed no differences in fungal richness among seeds from different plants (LMM; df = 2, F-value = 1.11, p-value = 0.37) (**Fig. 2e**), and among seeds from pods within the same plant (LMM, p-value > 0.05) (**Fig. 2f**). To summarize, these results suggest that seed bacterial/archaeal alpha diversity, but not fungal, varied plant to plant.

**Fig 2.**
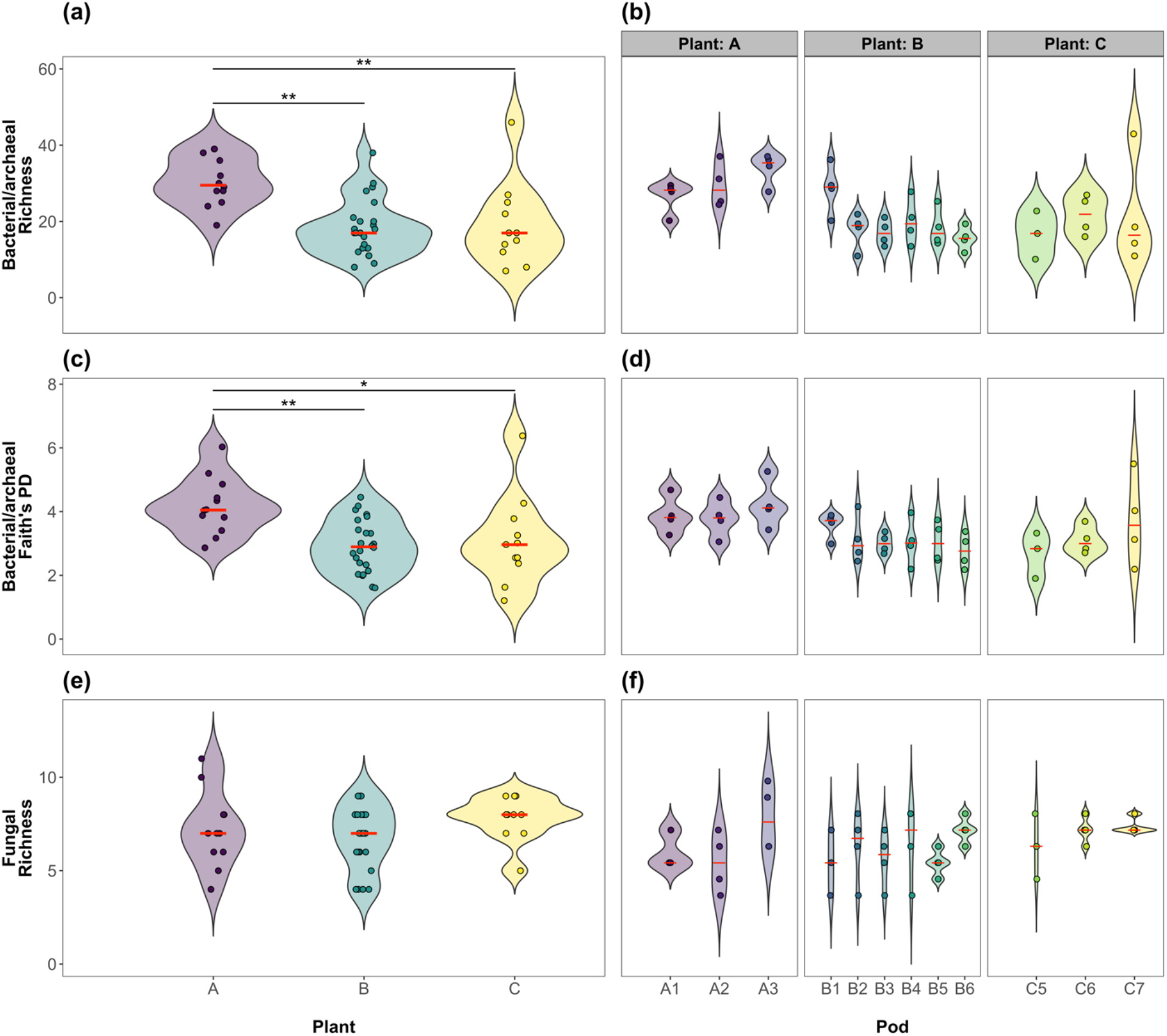
Bacterial/archaeal richness among plants were different (linear mixed-effects model, LMM; df = 2, F-value = 6.91, p-value = 0.015) (a), but not among pods within plant (p-value > 0.05) (b). Specifically, plant B and C displayed lower bacterial/archaeal richness compared to the plant A (Tukey’s HSD post hoc test; p-value = 0.001 and 0.006, respectively). Bacterial/archaeal phylogenetic diversity among plants were different (linear mixed-effects model, LMM; df = 2, F-value = 6.56, p-value = 0.003) (c), but not among pods within plant (p-value > 0.05) (d). Specifically, plant B and C displayed lower bacterial/archaeal phylogenetic diversity compared to the plant A (Tukey’s HSD post hoc test, p-value = 0.001 and 0.013, respectively). Fungal richness was not different among plants (linear mixed-effects model, LMM; df = 2, F-value = 1.11, p-value = 0.37) (e) and among pods within plant (p-value = 0.55) (f).

We detected a difference in seed bacterial/archaeal structure among plants (nested PERMANOVA, df =2, F-ratio = 2.94, p-value = 0.001) (**Fig. 3a**), but, again, not among pods from the same plant (nested PERMANOVA, df =9, F-ratio = 0.99, p-value = 0.63). Though separation among pods and plants are not obvious on the PCoA for the fungal seed microbiomes, we detected modest differences in fungal community structure among seeds from different plants (nested PERMANOVA, df =2, F-rati0 = 1.55, p-value = 0.02) (**Fig. 3b**), as well as among seeds from pods from the same plant (nested PERMANOVA, df =9, F-rati0 = 1.27, p-value = 0.03). An analysis of beta-dispersion revealed that there were differences in seed microbiome dispersion across different plants for bacterial/archaeal communities (PERMADISP, df = 2, F-value = 63.9, p-value = 0.001) (**Fig.3c**), but not for fungal communities (PERMADISP, df = 2, F-value = 0.22, p-value = 0.798) (**Fig. 3d**). Therefore, statistical differences in the seed microbiome across plants for the bacteria/archaea may be attributed to either centroid or dispersion, while fungal seed communities were different by centroid.

**Fig 3.**
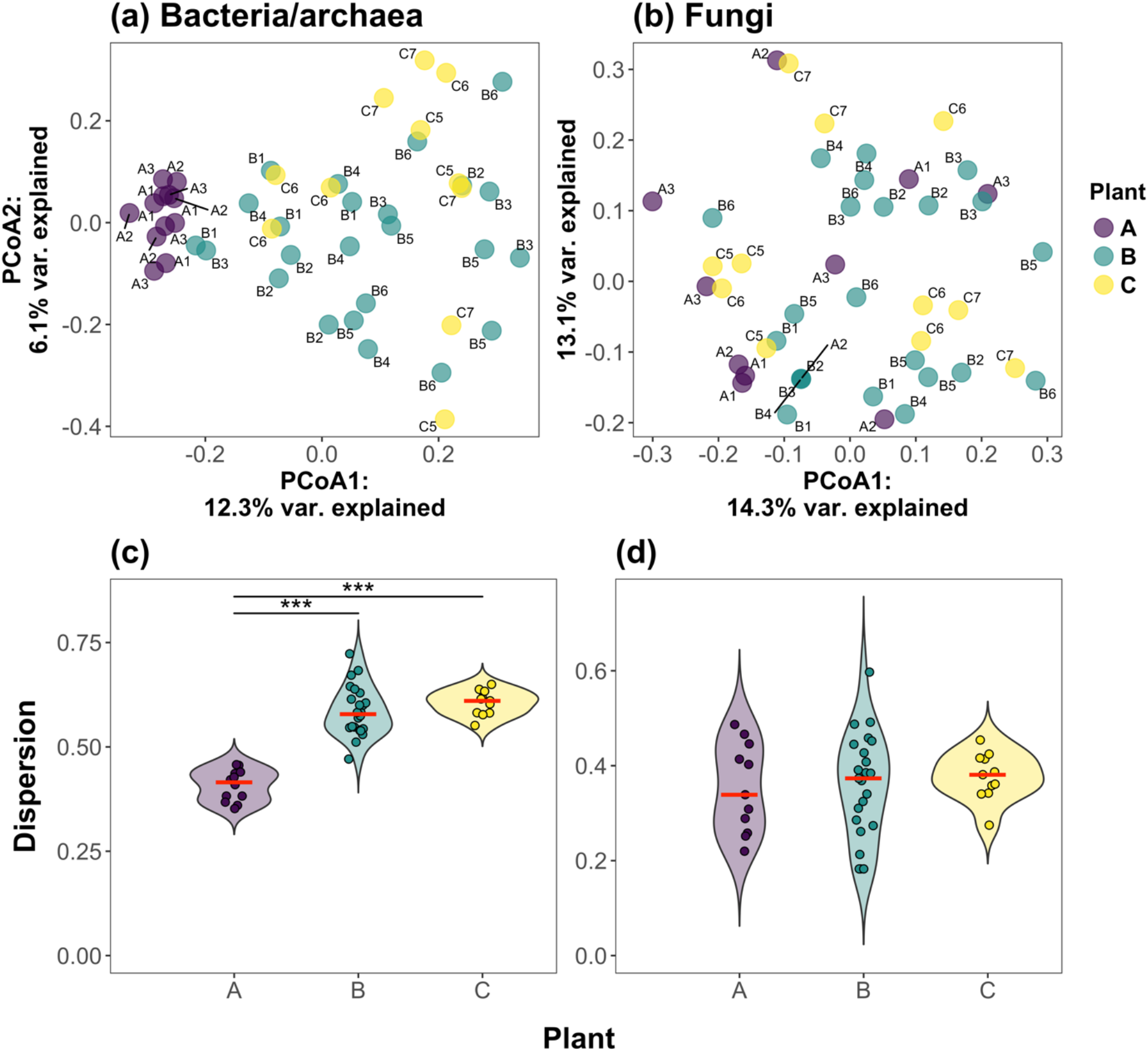
Principal coordinate analysis (PCoA) plot based on Jaccard dissimilarities of bacterial/archaeal (a) and fungal (b) OTUs. The samples were plotted and grouped based on plant as illustrated different colors. Each point was labelled by pod. Statistical analysis showed that seed bacterial/archaeal community structure differ among plants (nested PERMANOVA, df=2, F-ratio = 2.94, p-value = 0.002) but not pods (nested PERMANOVA, df =9, F-ratio = 0.99, p-value = 0.63). Statistical analysis also showed that seed fungal community structure differs among plants (nested PERMANOVA, df =2, F-rati0 = 1.55, p-value = 0.023) and pods (nested PERMANOVA, df =9, F-rati0 = 1.27, p-value = 0.03). Distance to centroid analysis using betadisper function from the vegan package revealed that there is variation in bacterial/archaeal Beta diversity among plant (PERMADISP, df = 2, F-value = 63.9, p-value = 9.6e-14) (c). In contrast, there were no variation in fungal Beta diversity among plant (PERMADISP, df = 2, F-value = 0.22, p-value = 0.802) (d).

### Bean seed microbiome composition

We identified 135 bacterial/archaeal and 49 fungal taxa at the genus level. The bacterial/archaeal individual seed communities were dominated by taxa from class Gammaproteobacteria (50.47%), Bacilli (24.48%), Alphaproteobacteria (8.68 %), and Bacteroidia (6.59 %) (**Fig. 4a**), and include Pseudomonas (13.58 %), Bacillus (10.2 %), Acinetobacter (9.5 %), Raoultella (7.09%), and Escherichia-Shigella (5.19%) as the major genera. Among members of the class Alphaproteobacteria, we also found genera Bradyrhizobium and Allorhizobium-Neorhizobium-Pararhizobium-Rhizobium with relative abundance of 2.57 and 0.85 %, respectively. Although seed fungal community composition varied among plants and also pods within plant, the fungal community was dominated by taxa belonging to classes Pezizomycetes (53.44 %), Agaricomycetes (25.7 %), and Dothideomycetes (11.17 %) (**Fig. 4b**), and the genera Helvella (53.44 %), Gautieria (19.65%), Acidomyces (7.29 %), Capnodiales_unidentified_sp_23791 (2.52 %), and Phlebiopsis (1.82%).

**Fig 4.**
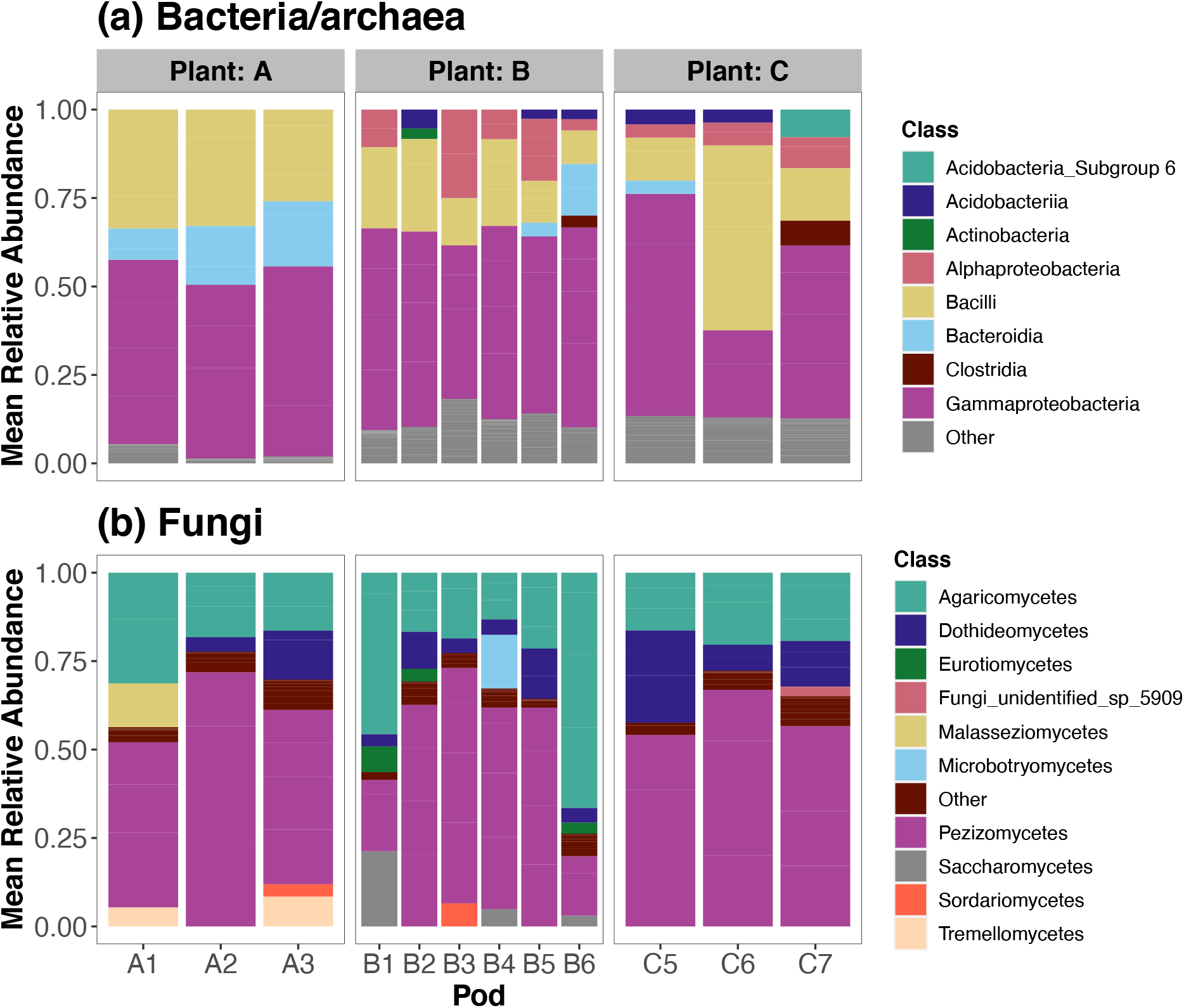
Bar plot represents mean relative abundances of bacterial/archaeal (a) and fungal (b) classes detected across plants. For bacteria/archaea, each pod consisted of 4 seeds (except for C5; 3 seeds); and for fungi, each pod consisted of 4 seeds (except for A1, B1 and C5; 3 seeds). The endophyte microbiome was assessed from the DNA extracted from single seed collected from each pod. Bacterial/archaeal and fungal classes with mean relative abundances of less than 10 % were grouped into the ‘Other’ classification, which includes many lineages (not monophyletic).

A key objective of this research was to understand the sources of variability in the individual bean seed microbiome to inform future study design. Because we found that the plant-to-plant seed microbiome variability was highest when grown in control conditions, we performed a power analysis to determine how many plants would be required to observe a treatment effect from seed samples pooled per plant. To detect the effect of treatment to bacterial/archaeal richness and phylogenetic diversity, pooled seeds from 9 and 12 plants are needed, respectively, for 16S rRNA richness and phylogenetic diversity, to achieve power of 0.8; and 13 and 19 plants to achieve power of 0.95 (**Fig. 5**).

**Fig 5.**
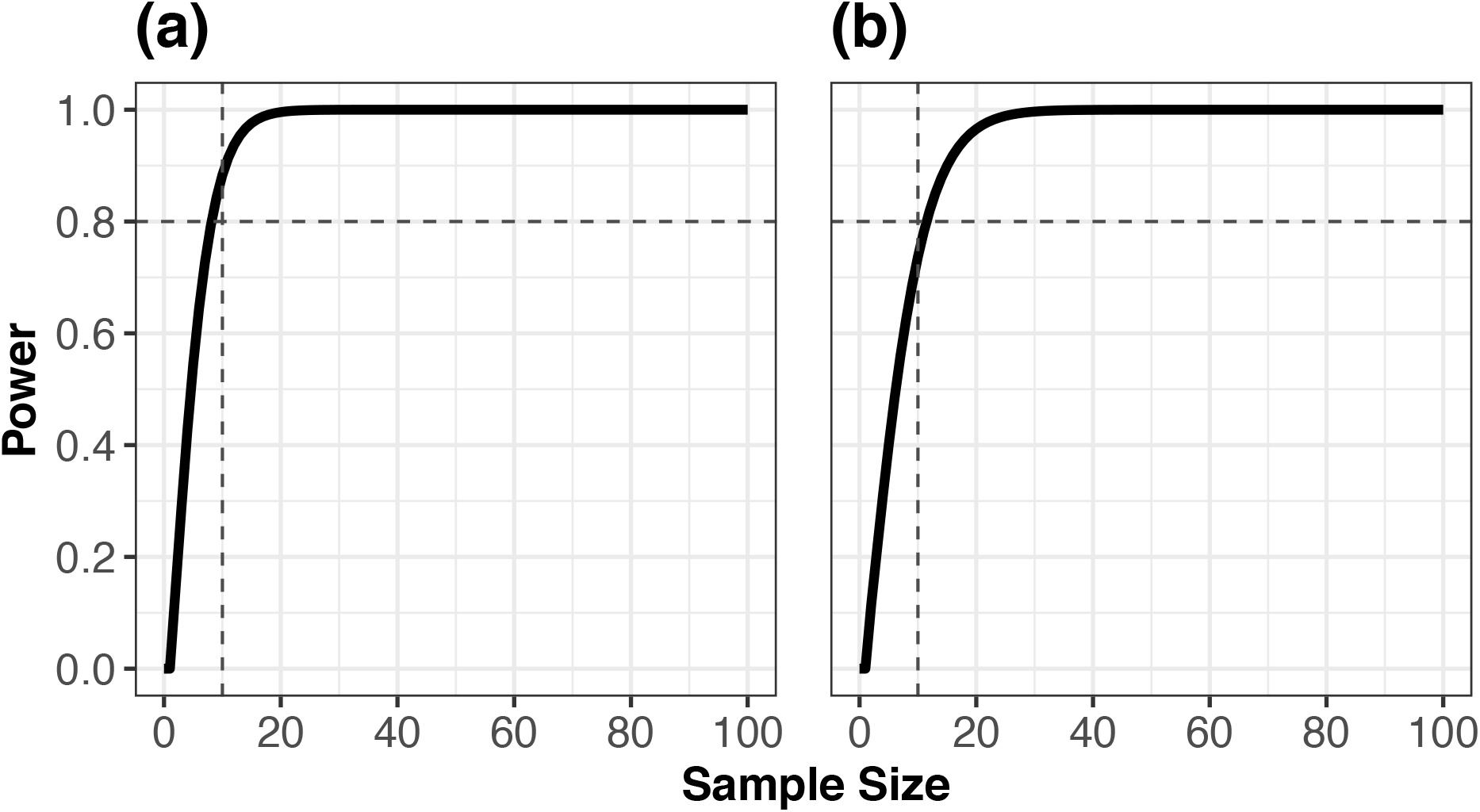
Analysis of power using pwr.t.test () function from the pwr package revealed that an effect of treatment on the 16S rRNA bacterial/archaeal alpha diversity (richness (a) and phylogenetic diversity (b)) would be detectable 12 plants at a power of 0.8. Because the highest seed microbiome variability was at the maternal plant level, individual seed microbiome sequence profiles were pooled in silico by plant to perform this power analysis at the individual plant level.

## Discussion

There remain gaps in our understanding of the persistence and assembly of seed microbiome members, especially across plant generations, and which microbiome members are beneficial and actively selected by, or even co-evolved with, the host. Here, we investigated the variability of the common bean microbiome at the resolution of the individual seed, which is the unit that delivers any vertically transmitted microbiome to the offspring. Because multiple legume seeds within a pod develop as a result of a single flower pollination, one simple hypothesis is that the individual seeds within a pod may harbor a highly similar microbiome if the floral pathway of assembly is prominent. However, recent work has suggested that the endophytic seed microbiome of green bean varieties of common bean likely colonize predominantly via the internal vascular pathway, and not the floral pathway (Chesneau et al. 2020), which may result in more homogeneity among seed microbiomes of the same plant. Our data support this finding, as seeds from the same plant (and therefore a common vascular pathway across pods) had relatively low microbiome variability, especially as compared across plants. It is expected that the vascular pathway of seed microbiome assembly is more likely to colonize the internal seed compartments (e.g., embryo), and therefore more likely to be vertically transmitted (Barret et al. 2016). It is yet unclear whether plant species that have a stronger relative importance of the floral pathway in seed microbiome assembly may exhibit higher microbiome variability at the pod/fruit level. Such an outcome may indicate that the experimental unit should instead be the pod level rather than the plant level for plant species dominated by floral assembly pathways.

There are many challenges in analyzing the microbiome of seeds generally and of a single seed in particular, which may be why cultivation-independent studies of single seeds are few (Abdelfattah et al. 2021). Previous studies showed that seeds have low microbial biomass and diversity (Adam et al. 2018; Chesneau et al. 2020; Rezki et al. 2016), especially relative to other plant compartments or soil. Therefore, many studies pool seeds to analyze the aggregated microbiome of many seeds and to get enough microbial biomass for metagenomic DNA extraction (Latz et al. 2021; Bergna et al. 2018; Wassermann et al. 2019; Adam et al. 2018; Johnston-Monje and Raizada 2011; Klaedtke et al. 2016). Generally, microbiome samples that have low biomass have numerous challenges in sequence-based analysis, as discussed elsewhere (Eisenhofer et al. 2019; Bender et al. 2018). First, unknown contaminants, either from nucleic acid kits or from mishandling of the samples, can have relatively high impact on the observed community composition, and so extraction and PCR controls are needed for assessment of contaminants and subtraction of suspected contaminants from the resulting community (Davis et al. 2018). Second, the sparse datasets (e.g., many zero observations for many taxa in many samples) generated from low biomass samples often require special statistical consideration and data normalization (Weiss et al. 2017; Anderson et al. 2011).

Plant host contamination of the microbiome sequence data is another consideration expected with analysis of the seed, and this challenge also applies to other plant compartments (Fitzpatrick et al. 2018; Song and Xie 2020). For 16S rRNA amplicon sequencing, the contaminant reads typically derive from host mitochondria and chloroplasts, but ITS or 18S amplicon analysis may also have reads annotated as Plantae. Therefore, nucleic acid extractions may attempt minimal disturbance of the plant tissue that is that target of microbiome investigation; for example, grinding tissues to include in the extraction will result in higher plant DNA contamination than separating microbial biomass from intact tissue. For seeds in particular, it is known that seeds can exude both antimicrobials and attractants to select for particular microbial members early in microbiome assembly of the germinated seed and emerging seedling (Chesneau et al. 2020; Meldau et al. 2012), and there is an active zone of plant and microbiome activity at the seed-soil-interface of a germinating seed (the spermosphere, e.g., Schiltz et al. 2015). Therefore, to target the native endophytic seed microbiome without also allowing for the plant’s potential selection for or filtering against particular members, it is important to use dormant seeds and also to minimally disrupt the seed compartment during extraction. Notably, many protocols have opted to first germinate seeds and, therefore, study the outcome of any plant selection prior to analyzing the seed microbiome (Wassermann et al. 2021; Bergna et al. 2018; López-López et al. 2010).

Taking all of these methodological aspects into consideration, this study presents a protocol and analysis pipeline for endophyte microbiome DNA extraction from a single dormant seed that experiences minimal tissue disruption in the extraction process, includes both positive and negative sequencing controls, and includes bioinformatic steps to identify contamination and remove host signal from the marker gene amplification. Notably, we chose to perform microbiome analysis based on a presence/absence taxon table rather than a table with relativized taxon abundances. This was done in consideration of the ecology of the seed endophyte microbiome members to likely be dormant until germination (Cope-Selby et al. 2017), and therefore the differences in relativized abundances do not reflect differences in fitness outcomes inside the dormant seed. We acknowledge that relative abundances could reflect differential microbiome member recruitment by the host plant, but this is not the objective of the study and would be best addressed with a different design to determine the multi-generation consistency and transmission rates of any observed enrichments, which would be supported by assessment of the seed microbiome within individual seeds, and across plant generations.

In conclusion, individual seed microbiome assessment provides improved precision in our understanding of plant microbiome assembly and sets the stage for studies of vertical transmission. We found that seeds produced by an individual bean plant can be considered as a unit (for comparative treatment study designs), and that seeds produced by different plants are expected to have slightly different microbiomes, even if grown under the same, controlled conditions and in the same soil source. Future work may consider whether functional redundancy in plant beneficial phenotypes across seed microbiome members may provide one mechanism for consistent outcomes in beneficial plant microbiome establishment.

## Acknowledgments

This research was supported by USDA 2019-67019-29305. AS acknowledges support from the USDA National Institute of Food and Agriculture and Michigan State University AgBioResearch (Hatch). AFB acknowledges a doctoral fellowship from the Fulbright Foundation and from Michigan State University. NS acknowledges support from the Michigan State University Plant Resilience Institute.

## Figure Legends

**Figure S1**. The proportion of plant reads of the total bacterial/archaeal (a) and fungal (b) reads showed that more than 90 % reads obtained were plant contaminants.

