## Supplementary Material for "Endophytic microbiome variation at the level of a single plant seed"

### Supplementary File: Background and Protocol for Microbial DNA extraction of an individual seed

By A. Fina Bintarti, Abby Sulesky-Grieb, Nejc Stopnisek and Ashley Shade

#### Background to seed microbiome assessment

Seed microbiome assessment has been conducted either by culture-dependent or culture-independent methods or a combination of both. Culture-dependent approaches are limited by technical difficulties in isolating microbes from seeds because not all members can be cultivated on agar plates. Seed microbiome members are assumed to be in a dormant stage until the plant germinates (Cope-Selby et al., 2017), and this may contribute to its difficulties to cultivate because they may need specific nutrients and growth conditions to be able to grow. Thus, culture-based methods often fail to detect all the microbial community members and can lead to biased results. Community profiling approaches using next-generation high-throughput sequencing of marker genes, such as 16S rRNA and ITS genes for bacteria/archaea and fungi may provide a better assessment of microbial community in the seed and a more comprehensive picture of microbial community structure. A number of studies have shown variability of the microbial community in the seed from different plant species and genotypes (Özkurt et al., 2020; Wassermann et al., 2019), different geographical sites (Chartrel et al., 2021), or even between seed developmental stages (Barret et al., 2015) and seed compartments (Eyre et al., 2019).

To our knowledge, this is the first study to use marker gene high-throughput sequencing methods to assess the microbial community of individual seeds of common bean (*Phaseolus* *vulgaris*) to investigate its variability among plants and pods. Understanding on seed-to-seed, pod-to-pod, and plant-to-plant microbiome variability provides essential information on pooling biological samples and allows well-powered experimental design on the seed microbiome assessment under plant treatments. Extracting metagenomic microbial DNA from one seed is extremely difficult because the individual seed is considered as a low-microbial-biomass sample, and microbial DNA extraction from low-microbial-biomass samples can be a major challenge in studying the microbial community and ecology. Hence, a robust and efficient DNA extraction from low microbial samples is a crucial step because reproducibility and accuracy of microbiome study with amplicon-based sequencing approaches will depend on the efficient DNA extraction from the sample (Davis et al., 2019).

Moreover, microbial DNA extraction from low-microbial-biomass and low diversity samples is prone to DNA contaminations from other microbes and/or plant contaminants, such as mitochondria and chloroplasts. Thus, it is necessary to set up strategies to minimize DNA contamination during extraction as well as in the downstream analysis. We developed and optimized protocols for microbial DNA extraction from individual seed samples of common bean (*P. vulgaris*) var. Red Hawk. The protocols described in this study were aimed to generate robust methods that can be generally implemented to study seed microbiomes.

**Challenges to seed microbiome DNA extraction protocols**

Total DNA extraction from one seed sample can be problematic and challenging. There are some key limitations that we need to consider and carefully assess before conducting DNA extraction from seed samples.

**1.) Low diversity.** Previous studies show that seeds have low microbial diversity (Adam et al., 2018; Chesneau et al., 2020; Rezki et al., 2016) relative to other plant compartments, or rhizosphere and soil. Since seeds have low microbial diversity, it is important to include a mock microbial community as a positive control for assessing PCR amplification and sequencing efficiency. Since the expected composition of the mock microbial community is known, we can estimate any sequencing error (e.g., chimera), identify diversity biases, and determine microbial contaminations by including the mock community in a sequencing run (Pollock et al., 2018).

**2.) Low microbial biomass.** Seeds, particularly individual seeds, contain low microbial biomass compared to soil or rhizosphere samples. One of the challenges in working with low microbial biomass samples (low DNA target) is the feasibility and efficiency of the DNA extraction method and exogenous microbial contamination. Our strategy to overcome this issue is to include whole-cell mock microbial community as the DNA extraction positive control to establish the DNA extraction procedure. We also include a negative control (buffer only) that is important for assessing microbial contamination in the samples.

**3.) Plant anti-microbial chemicals (plant defense compounds).** One hypothesis on why seeds have low microbial diversity is the occurrence of population bottlenecks in the seed (Newcombe et al., 2018), especially individual seeds, that is caused by the accumulation of anti-microbial compounds in the seed (Chesneau et al., 2020; Meldau et al., 2012). These compounds are activated when seeds are crushed or germinated. Thus, we performed buffer soaking methods instead of grinding the samples or germinating the seeds for the DNA extraction procedure to avoid anti-microbial compounds affecting our results.

**4.) Plant contamination.** Plant contamination in plant microbiome study is common because plant compartments including the seed contain plastids, chloroplasts, and mitochondria that share common ancestry and have sequence similarity with bacteria. There are three main approaches to minimize plant host contamination including modification of microbial DNA extraction to prevent the co-extraction of plant organelles, the application of PCR amplification blocking primers to block the amplification of plant sequences, and the use of specific mismatch primers (Beckers et al., 2016). (Lundberg et al., 2013) constructed PNA clamps for plastids (pPNA) and mitochondria (mPNA) that can bind tightly to the contaminant sequences and block its amplification. Another approach is the use of anti-chloroplast primers 799F that can amplify 16S rRNA gene sequences but also avoid the amplification of plant sequences. Beckers et al. (2016) reported that primer pair 799F/1391R was the most efficient in eliminating plant DNA (very low amplification of plant DNA) and resulted in the highest amount of bacterial OTUs.

However, there is a scientific motivation to be able to directly compare microbiome data across studies (for instance, to compare with other studies that include soils, plants, potential sources of dispersal/immigration). Thus, the use of the popular Earth Microbiome Project 16S rRNA V45 primers is often desirable (https://earthmicrobiome.org/), despite that these primers co-amplify plant contaminants. Therefore, other steps to reduce host signal can be taken in the DNA extraction protocol. Specific to this seed microbiome study, we performed an adaptation of microbial DNA extraction to prevent the co-extraction of plant organelles. Instead of grinding the seed that can release plant organelles, we used a phosphate buffered saline (PBS) soaking procedure. This procedure has been used by previous studies in assessing seed microbiomes

(Barret et al., 2015; Rezki et al., 2016; Torres-Cortés et al., 2018). By using a seed soaking procedure, microbial cells in the seed coat and funiculus will be released to the suspension (Gitaitis & Walcott, 2007; Rezki et al., 2016).

**5.) Non-host DNA contamination.** As we described above, DNA contamination introduced during the DNA extraction method is a major challenge in assessing microbial communities from low microbial biomass samples. There are different strategies in removing DNA contaminants before and after sequencing. In this study, we included blank or negative controls for the DNA extraction method as well as for PCR amplification. Strategies can also be performed after sequencing, for example, by removing any microbial taxa that have been published and identified as a common contaminant. However, this method cannot be implemented for all studies because the observed microbial community is different for each study. Another method is removing taxa that are also present in the negative control. However, this strategy may also remove the actual members of the microbial community because of multiplexing artifacts that occur in the negative control (Karstens et al., 2019). In our study we performed decontamination using an open-source R package called microDecon for identifying and removing contamination (McKnight et al., 2019).

**Protocol for Cultivation-Independent Native Seed Endophyte Analysis**

We performed surface sterilization of the seed samples before extracting the DNA because our study focused on the seed endophytic communities. Surface sterilization of sample is a required procedure to study plant endophytes (Saldierna Guzmán et al., 2020) because we need to completely remove the epiphytic microbes from the seed surfaces. The seed epiphytes are mostly derived from plant surfaces (e.g., leaves, stems, fruits) and/or environment (e.g., soil) (Shade et al., 2017). Surface sterilization also removes microbial contamination from human contact during harvesting, handling, and processing.

**Part 1. Seed surface sterilization and overnight soaking procedures**

**Expected time:** 20 minutes, overnight

**Materials**

1. Common bean seed (*P. vulgaris L.*, var. Redhawk) (approximately 0.6 g per seed)
2. Trypticase Soy Agar (TSA) and Potato Dextrose Agar (PDA) plates
3. Sterilization solution: 10 % (v/v) bleach with 0.1% (v/v) Tween20
4. Sterile Phosphate Buffer Saline (PBS) 1 X with 0.05 % (v/v) Tween20

**Equipment**

1. 50 mL centrifuge tube (USA Scientific, VWR)
2. Beaker
3. Analytical balance
4. Sterile dissecting scalpel (size 20)
5. Sterile dissecting forceps
6. Sterile disposable Petri dishes
7. Orbital shaker
8. Plate spreader or plating beads

**Procedure**

1.) Select healthy seeds with no disease symptoms from the stock and weigh the seeds to obtain seed mass data.

2.) Place seed(s) into a sterile 50 mL centrifuge tube and immerse the seed in ~ 20-25 mL sterilization solution (10 % (v/v) bleach with 0.1% (v/v) Tween20) for 10 minutes.

- *A different volume of sterilization solution can be used, based on the number/size of seeds.*
- *Shake the tube several time during incubation.*

3.) Discard the sterilization solution and rinse/wash the seed with sterile water 5 times to remove bleach residue.

- *To check the effectiveness of surface sterilization, spread 50-100 µl of the final rinse water on to TSA and PDA plates. Incubate the TSA and PDA plates at 30 ˚C for 2-3 days and 25-26 ˚C for 5 days, respectively. Discard associated sample if there is any microbial growth on the plates.*

4.) Place sterile seed onto sterile plate and carefully dissect/open the seed in half long-ways on the natural division of cotyledon using sterile surgical blade and forceps.

- *In this study, we removed the seed coat instead of dissecting the seed in half. The purpose of seed coat removal is because our study focused on seed endophytes, we assumed that removal the seed coat could increase the release of the endophytes located in the endosperm and embryo into the buffer solution. However, we observed high plant contamination in when the seed coat was removed (more than 90% of total reads). We also found that removing the seed coat is time consuming and produces plant debris that can interfere with the DNA extraction process and can be the source of chloroplast and mitochondria contamination. Thus, we propose to dissect/open the seed in half long-ways on the natural division of cotyledon instead of removing the seed coat. In our experience, this allows for the release of endophytes into the buffer and minimizes host contamination from seed coat removal.*

5.) Immerse and soak surface sterilized seed in sterile Phosphate Buffered Saline (PBS) 1X supplemented with 0.05% (v/v) Tween 20 (3 mL) under constant agitation (160 rpm) overnight at 4 ˚C.

- *A different volume of buffer can be used based on the number/size of seed sample.*
- *We recommend to always include a DNA extraction positive control for low microbial biomass samples like seeds (e.g., a mock microbial community). We used the commercial ZymoBIOMICS Microbial Community Standard (catalog number: D6300) for this study by diluting 75 µL (1 prep) of the mock community into 3-5 mL PBS 1X with 0.05% (v/v) Tween20. Also, we created our own mock community in-house to include particular bacteria and fungi that reflect the expected composition of common seed microbial community members. The mock community included populations of type strains or isolates grown in the lab, and then combined at an equal ratio at a concentration of $10^8$ cells per ml for bacteria ($10^6$ cells per ml for Streptomyces) and $10^7$ cells per ml for fungi and stored in glycerol stock in the -80 ˚C. Therefore, the positive control DNA extraction of our in-house mock-community would be performed directly on these cells and can be sequenced and checked for contamination from the expected composition.*
- *We recommend to always include a DNA extraction negative control of extraction buffer only (3-5 mL PBS 1X with 0.05% (v/v) Tween20). This sample should be sequenced to check for contamination and to calculate a sequencing error rate (Schloss et al., 2011).*

**Part 2: Seed processing and pellet collection**

**Expected time:** 90 minutes

**Stopping points:** It is recommended to either stop after the pellet collection step, or to go through the DNA extraction protocol in the same day

**Materials**

1. Overnight-soaked seed in sterile PBS 1X with 0.05%. (v/v) Tween20

**Equipment**

1. Swinging-bucket rotor centrifuge
2. Vortex
3. Sterile forceps
4. Beaker
5. Micropipette
6. Sterile barrier micropipette tips
7. Microcentrifuge tubes

**Procedure**

6.) Centrifuge all samples and controls at 4500 x g for 60 minutes at 4 ˚C.

- *We used a centrifuge with swinging-bucket rotor rather than fixed-angle rotor so that the pellets will form at the bottom of the conical tube, thus, it is easier to resuspend and collect the pellets. The original protocol from previous study (Barret et al., 2015) stated that centrifugation was performed at 6000 x g for 10 minutes at 4 ˚C. Since the maximum speed for swinging-bucket rotor centrifuge is 4500 x g, we extended the centrifugation time.*

7.) Carefully remove seeds aseptically with sterile forceps, spin tubes again with bucket centrifugation at 4500 x g for 10 min at 4 ˚C to re-pellet any disturbed material. Carefully remove supernatant with sterile disposable pipette or micropipette until ~1-2 mL remain.

- *Alternatively: After first hour of centrifugation, gently pour most of the supernatant out of the tubes and discard, then aseptically remove seeds with sterile forceps, leaving approx. 1-2 mL of supernatant in the tube.*

8.) Resuspend pellet in remaining supernatant by vortexing for ~1 minute.

9.) Transfer the suspension into 1.5 or 2 mL microcentrifuge tube and centrifuge at 20,000 x g for 10 minutes.

10.) Discard the supernatant and keep the pellet for DNA extraction using E.Z.N.A.®Bacterial DNA Kit with centrifugation protocol.

- *Pellets can be stored at -20˚C until they are ready to be extracted.*

**Part 3: Microbial DNA extraction from seed pellet with bead beating procedure using E.Z.N.A.® Bacterial DNA Kit with modification**

**Expected time:** 4 hours active time, 3 hours of incubation time

**Materials**

1. Seed pellet collected from the previous step
2. E.Z.N.A.® Bacterial DNA Kit (D3350-02) (OMEGA Bio-Tek Inc., Norcross, GA, USA)
3. 100 % Ethanol
4. Tris-EDTA (TE) Buffer, Molecular Biology Grade (pH 8.0)

**Equipment**

1. Micropipette
2. Sterile barrier micropipette tips
3. Microcentrifuge tubes
4. Vortex
5. Beaker
6. Heat block or water bath

Before starting:

- Prepare HBC Buffer, DNA Wash Buffer, and Lysozyme kit components as instructed in the manufacturer's protocol
- Set a heatblock or water bath at 37 ˚C
- Set a shaking heatblock or water bath at 55 ˚C
- Set an incubator or a heatblock at 65 ˚C (can change the 37 ˚C to 65 ˚C later in the protocol, if necessary)
- Heat Elution Buffer to 65 ˚C

**Procedure**

11.) Add 100 µL TE buffer to the pellet and completely resuspend the pellet.

12.) Add 10 µL Lysozyme resuspended with Elution Buffer (see bottle for instructions). Vortex to mix thoroughly. Incubate in 35 ˚C heat block for 1 hour.

- *We used 1 hour incubation instead of 10 minutes as stated on the manufacturer's protocol to achieve complete digestion of the cell wall.*

13.) While incubating, aseptically add 25 mg Glass Beads S (included with the kit) to new, labeled, 1.5 mL tubes.

14.) After incubation transfer entire sample, including any material that has precipitated out, into the corresponding tube with glass beads.

15.) Vortex the bead-beating tubes at maximum speed for 10 minutes. After vortexing, allow tubes to rest a few minutes for glass beads to settle out. Transfer supernatant to clean 1.5 mL tube.

- *We implemented a bead-beating step into the protocol for hard-to-lyse bacteria/archaea and fungi. This procedure yielded better results (higher DNA concentration) than extractioins without a bead-beating step.*
- *We extended the vortexing time at maximum speed from 5 minutes to 10 minutes for optimal cell wall disruption.*

16.) Add 100 µL TL Buffer and 20 µL Proteinase K Solution to all tubes. Pipette up and down to break up pellet, if present, and then vortex to mix thoroughly. Incubate at 55 ˚C in a shaking heat block for 2 hours (500 rpm). Alternatively, incubate in a stationary heat block and vortex every 20 minutes.

- *We used longer incubation time for optimal DNA yield.*

17.) Add 5 µL RNase A. Invert tube several times to mix. Let sit at room temperature for 5 minutes.

18.) Centrifuge at 10,000 x g for 2 minutes to pellet any undigested material.

19.) Transfer the supernatant to a new 1.5 mL microcentrifuge tube. Do not disturb the pellet. Discard pellet.

20.) Add 220 µL BL Buffer. Vortex to mix. Incubate at 65 ˚C for 10 minutes. (after this step, aliquot the needed amount of elution buffer into a tube and place in the 65˚C block to warm for later use).

21.) Add 220 µL 100% ethanol. Vortex for 20 seconds at maximum speed to mix thoroughly. Break any precipitates by pipetting up and down 10 times.

22.) Insert a HiBind® DNA Mini Column into a 2 mL Collection Tube. Transfer the entire sample to the HiBind® DNA Mini Column, including any precipitate that may have formed.

23.) Centrifuge at 10,000 x g for 1 minute. Discard the filtrate and the collection tube.

24.) Insert the HiBind® DNA Mini Column into a new 2 mL Collection Tube.

25.) Add 500 µL HBC Buffer diluted with 100 % isopropanol (see the bottle for instructions). Centrifuge at 10,000 x g for 1 minute. Discard the filtrate and reuse the collection tube.

26.) Add 700 µL DNA Wash Buffer diluted with 100 % ethanol (see the bottle for instructions). Centrifuge at 10,000 x g for 1 minute. Discard the filtrate and reuse the collection tube.

27.) Repeat Step #26 for a second DNA Wash Buffer wash step.

28.) Centrifuge the empty HiBind® DNA Mini Column at maximum speed (> 10,000 x g) for 2 minutes to dry the column.

- *We used a centrifuge with maximum speed of 20,000 x g for optimal removal of trace ethanol.*

29.) Insert the HiBind® DNA Mini Column into a new, nuclease-free 1.5 mL microcentrifuge tube.

30.) Add 30 µL Elution Buffer heated to 65 ˚C to the center of the HiBind® matrix. Let sit for 10 minutes at room temperature.

- *We decreased Elution Buffer volume from 50-100 µL as stated on the manufacturer's protocol to 30 µL to increase DNA concentration.*

- *To obtain more yield, second elution can be conducted with the same Elution Buffer volume.*

31.) Centrifuge at 10,000 x g for 1 minute to elute the DNA. Store the DNA at -20 ˚C for temporary storing or -80 ˚C for long-term storing.

- *We measured the DNA concentration using Qubit™ dsDNA HS (High Sensitivity) Assay Kit with the Qubit Fluorometer. This protocol yielded DNA with the concentration of ~0.7-1 ng per µL per gram of seed. Moreover, the PCR amplification of bacterial 16S V4 and fungal ITS2 also resulted in clear and specific bands.*
- *We tried the Qiagen DNeasy PowerSoil DNA Isolation Kit for the DNA extraction after collecting the seed pellets. However, the protocol using this kit was irreproducible. The DNA concentration was too low to be detected on the Qubit Fluorometer and the PCR amplification of bacterial 16S V4 and fungal ITS2 resulted in very weak or no specific bands. We assumed that the Qiagen DNeasy PowerSoil DNA Isolation Kit was not reliable enough to extract DNA from low microbial biomass samples, such as seeds or individual seeds, in particular. Thus, we chose to use the DNA extraction kit with optimum lysis that implement both enzymatic digestion of cell wall and physical disruption using bead-beating step.*

**Figure S1.** The proportion of plant reads of the total bacterial/archaeal (a) and fungal (b) reads showed that more than 90 % reads obtained were plant contaminants.

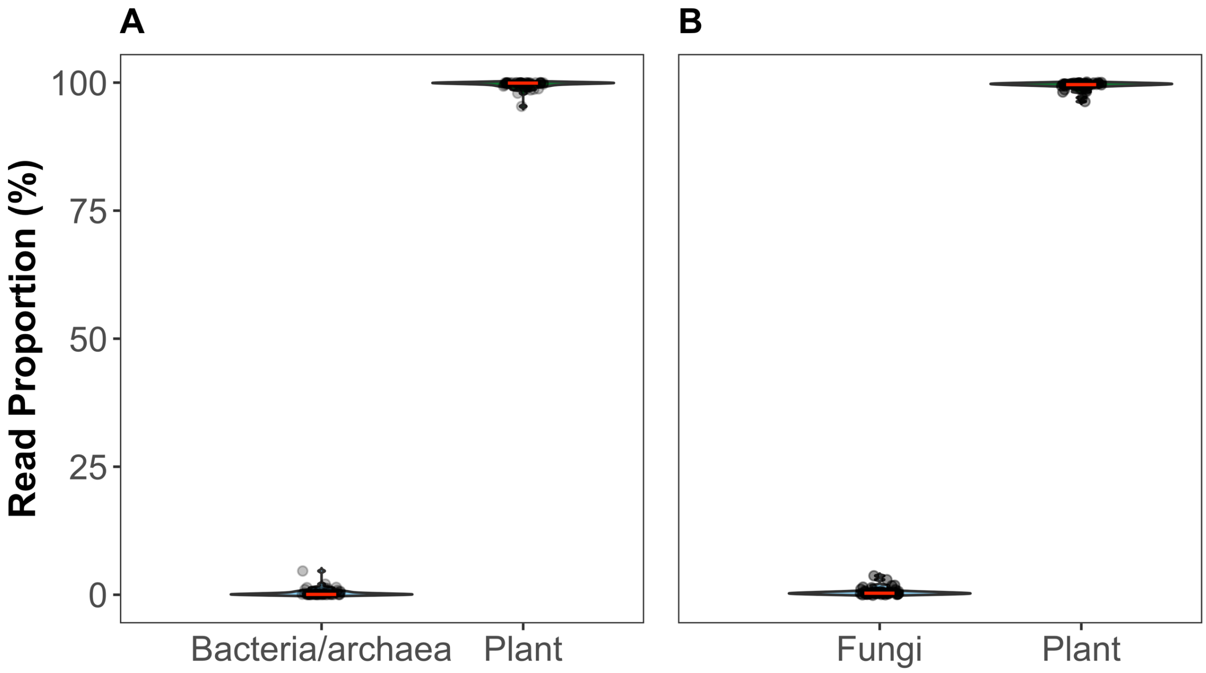
